# Mutagenic distinction between the receptor-binding and fusion subunits of the SARS-CoV-2 spike glycoprotein and its upshot

**DOI:** 10.1101/2021.11.15.468283

**Authors:** Robert Penner

## Abstract

We observe that a residue R of the spike glycoprotein of SARS-CoV-2 which has mutated in one or more of the current Variants of Concern or Interest or under Monitoring rarely participates in a backbone hydrogen bond if R lies in the S_1_ subunit and usually participates in one if R lies in the S_2_ subunit. A partial explanation for this based upon free energy is explored as a potentially general principle in the mutagenesis of viral glycoproteins. This observation could help target future vaccine cargos for the evolving coronavirus as well as more generally. A study of the Delta and Omicron variants suggests that Delta was an energetically necessary intermediary in the evolution from Wuhan-Hu-1 to Omicron.

## 1. Introduction

This short note isolates a specific and elementary observation about Protein Data Bank (PDB) [1] files concerning the mutated residues in the current Variants of Concern and of Interest plus the Variants under Monitoring, as per [15] on 22Oct2021, of the SARS-CoV-2 spike glycoprotein S. This observation is then applied to the appearance of the Omicron variant. It has not, to our knowledge, appeared in the literature other than in our own earlier work [19] in the context of specific Variants of Concern, and it may be material going forward in designing mRNA or other types of vaccine cargos, if necessary, as the coronavirus continues to evolve. It is anyway worth considering, in this by-now highly studied example, as a potentially more general example of viral glycoprotein mutagenesis, since we provide a partial explanation for our observation based upon general principles involving free energy.

To state this fact about mutagenic pressure in the spike, recall [4] that as in many examples of viral glycoproteins, in particular commonly with Class I fusion mechanisms [10,24], S is composed of two subunits S_1_ and S_2_, where S_1_ mediates receptor-binding extracellularly, and S_2_ mediates fusion within an endosome. One particularity of S is a host-furin mediated cleavage between S_1_ and S_2_, at residue number 685-686. There is furthermore a second cleavage lying adjacent to the fusion peptide, mediated by host-cathepsin or serine protease, of a C-terminal segment 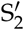;of S_2_, at residue number 815-816. See [2] for more information on cleavage in the SARS-CoV-2 spike.

In any protein, hydrogen bonds form between backbone Nitrogen atoms N_*i*_-H_*i*_ and Oxygen atoms O_*j*_=C_*j*_ in different peptide units, and these are called Backbone Hydrogen Bonds (or BHBs). (To be precise: A DSSP [13] hydrogen bond is accepted as a BHB provided that furthermore the distance between H_*i*_ and O_*j*_ is less than 2.7 Å and ∠*NHO* and ∠*COH* each exceed 90°.) A protein residue R_*i*_ itself is said to *participate* in a BHB if either the nearby Nitrogen N_*i*_-H_*i*_ donates or the nearby Oxygen O_*i*_=C_*i*_ accepts a BHB. (Again to be precise, if at least two monomers of the trimeric spike participate, then the residue itself participates.) On average for all proteins, roughly 70-80% of all residues participate in a BHB [7].

Here is the main easily confirmed empirical observation of this paper, which is quantified in Table 1 and subsequently discussed: A residue R of the SARS-CoV-2 spike glycoprotein S which has mutated in one or more of the current Variants of Concern or Interest or under Monitoring [15] (cf. Table 1 for these mutagenic residue numbers), rarely participates in a BHB if R lies in S_1_ and usually participates in a BHB if R lies in S_2_.

**Table 1.** **Summary Statistics of Variant Mutagenic Residues** for the PDB-covered submolecule S of the SARS-CoV-2 spike glycoprotein (residues 27 to 1147) and its sub-molecules S_1_ (residues 27 to 681) and S_2_ (residues 682 to 1147). Averages are per residue in each molecule summed over the 15 structures in the third and fourth columns. A residue is *missing* if it is not modeled in the PDB file (interpreted as disorganized), it is *absent* if it occurs in the PDB file but does not participate in either nearby backbone hydrogen bond (along the backbone), and it is *unbonded* if it is either missing or absent in at least 10 of the 15 PDB files in the database for that residue. In each case, the bar over the molecule denotes the subset of mutated residues M of Wuhan-Hu-1 among the Variants of Concern or Interest and those under Monitoring, namely, residues (5 9 12 18-20 26) 52 67 69-70 75-76 80 95 136 138 144-145 152 156-158 190 215 243-244 246-253 346 417 449 452 478 484 490 501 570 614 641 655 677 679 681 701 716 796 859 888 899 950 982 1027 1071 1092 1101 1118 (1176), where the residues in parentheses are outside PDB-coverage and are not reflected in this table. A residue contributes to the average free energy BFE only if it is bonded.

| Mol | #Res | avg #missing | avg #absent | avg #unbonded | avg BFE |
| --- | --- | --- | --- | --- | --- |
| S | 1121 | 1.39 | 4.91 | 0.35 | 2.41 |
| $\bar{S}$ | 56 | 4.71 | 5.34 | 0.61 | 2.47 |
| S<sub>1</sub> | 655 | 1.85 | 5.52 | 0.42 | 3.16 |
| $\bar{S}_1$ | 43 | 6.14 | 5.69 | 0.74 | 2.80 |
| S<sub>2</sub> | 466 | 0.73 | 4.05 | 0.25 | 1.60 |
| $\bar{S}_2$ | 13 | 0.00 | 4.15 | 0.15 | 2.14 |

Note that the missing and absent columns in the table come directly from the PDB and DSSP with no provisos (other than those conventions in parentheses in the text). The unbonded column presents our novel insights and depends upon a cutoff 5, namely, it is bonded (i.e., neither absent nor missing) in at least 5 of the 15 structures. The last BFE column depends not only on the cutoff 5 but also on our theory of BHBs. All of the preceding trends are invariant under changing this cutoff by unity, with this data not presented.

A general but not entirely satisfactory explanation for this involves the free energy of structural details stabilized by BHBs. Namely, viral glycoproteins which mediate receptor binding and membrane fusion are by their very nature metastable. It follows that successful viral mutation can neither increase free energy by so much as to disturb stability of the molecule nor decrease it by so much as to interrupt near-instability, for otherwise the molecule will respectively either explode or fail to reconform and function correctly. The minimal way to avoid this twofold constraint is to mutate residues that do not participate in BHBs at all, and that is precisely what we find in S_1_ before mutation. However, we shall also discover, most interestingly, that this is not reversible.

We shall discuss S_2_ subsequently, only after including certain salient definitions, facts, and data, and note in this Introduction simply that the existence of BHBs and their free energies are obviously functions of pH. This alone might account for differences between S_1_ and S_2_, since the endocytotic pathway is highly acidifying [4].

## 2. Materials and Methods

As is customary, we record mutations relative to an original Wuhan genome called Wuhan-Hu-1 and its corresponding spike protein (UniProt Code P0DTC2 [22]) by considering only structure files with resolution below some bound, for our purposes resolution at most 3.0 Å, neither cleaved nor bound to antibody or receptor, and computed via cryoelectron microscopy. These 15 exemplar structures 6VXX 6X29 6X79 6XLU 6XM0 6XM3 6XM4 6ZB5 6ZGE 7A4N 7AD1 7DDD 7DF3 7DWY 7JWY for S from the PDB depend upon various techniques of stabilizing S in its prefusion conformation [11,12,14,21,23,25–29]. The molecules are therefore not truly identical, hence the utility of taking consensus and average data across the collection of PDB files, as we shall do.

Some of the previous considerations can be calibrated by employing a new concept and quantity in structural biology, the so-called backbone free energy (BFE) from [17], which can be computed from a PDB file, to be called simply a *structure*. Roughly, the BFE of a structure stabilized by a BHB is computed from geometry [16] by comparing the planes containing the peptide units of the donor and accepter of the BHB, and applying the Pohl-Finkelstein quasi Boltzmann Ansatz [5,6,20].

Let us next briefly give a more complete discussion of the method from first principles, referring the interested reader to [16] for the background and data for general proteins (as explained presently), [17] for application to viral glycoproteins, [18] for application to coronavirus spikes, and [19] for the SARS-CoV-2 spike S in particular. One starts by choosing a suitably unbiased subset of the PDB and computing all of its attendant BHBs, comprising a collection of 1166165 BHBs for the unbiased subset in [16] and for us throughout. For each of these BHBs, there is a rotation of space from the peptide plane of its donor to the peptide plane of its receptor mapping the peptide bond of the former to that of the latter. This defines an a priori distribution on the space of all rotations, which is computed once and for all. Now given another BHB, there is an associated free energy given by taking the density of this a priori distribution at this new subject BHB, suitably normalized to give the BFE in kcal/mole. Thus, the geometry of the backbone described in a PDB file determines a BFE associated to each BHB in any protein. The fundamental fact, established in [17] for viral glycoproteins, is that residues of large BFE target locations of large conformational change in the backbone, in particular typically including the fusion peptide.

There is a trichotomy of possibilities for a residue R in a specific structure: R may be modeled in the structure and participate in a BHB or not, and in this latter case we say R is *absent*, but R may also simply be *missing* from the PDB file. (As before, these properties of residues are taken as consensus data from the three monomers.) R can be missing for a number of simple reasons: the protein may be disordered at R [4]; the experiment may be inaccurate or problematic at R; the data and its refinement may not model R to within reasonable parameters; or R may be C-terminal or N-terminal to the experimentally synthesized sub-peptide of the protein S.

The average of resolutions in our collection of structures is 2.77 Å, and of percentage Ramachandran outliers is 0.1, so these are all high-quality experimental structures. (Note that Clashscores and Sidechain Outliers are not particularly relevant measures of quality for our purposes.) As argued in [19], it follows that the first among the possibilities for R missing is the most likely, so one might *conflate missing with disordered* for high quality structures within PDB-range. The consensus range of our collection of structures for S is comprised of residue numbers 27 to 1147.

A residue that is missing or absent is said to be *unbonded* and is *bonded* otherwise. If a residue R_*i*_ is bonded, then it participates in a BHB so there is either a BHB with donor N_*i*_-H_*i*_ or one with acceptor O_*i*_=C_*i*_ or both, and the BFE of the residue R_*i*_ is defined to be the maximum of the BFEs of these one or two BHBs, first averaged over the two or three monomers. If a residue is unbonded, then its BFE is undefined.

Specifically to give a quantitative sense to what follows, the range of BFE values is -2.9 to +6.85 kcal/mole with approximate 50th, 90th, and 99th percentile cutoffs respectively given by 1.4, 4.6, and 6.6. The validated hypothesis is that if the BFE of a residue lies in the 90th percentile, i.e., is at least 4.6 kcal/mole, then within one residue of it along the backbone, the sum of the two adjacent backbone conformational angles changes by at least 180 degrees in its pre- to post-fusion reconformation. The converse does not hold.

## 3. Results

### 3.1. Wuhan-Hu-1 to Many Variants

As argued before in order to preserve metastability of the molecule, the BFE before and after a mutation must be more or less constant across the spike, and the plot of BFE across the spike is depicted in Figure 1. Higher BFE is evidently concentrated in S_1_ compared to S_2_. The several regions of meaningful negative BFE are illustrated in the figure by the intersections of the plot with the grey horizontal line, which correspond to nearly ideal *α* helices. Notice that each cleavage site is surrounded by a region of high BFE, and likewise for the two ends of HR1.

**Figure 1.**
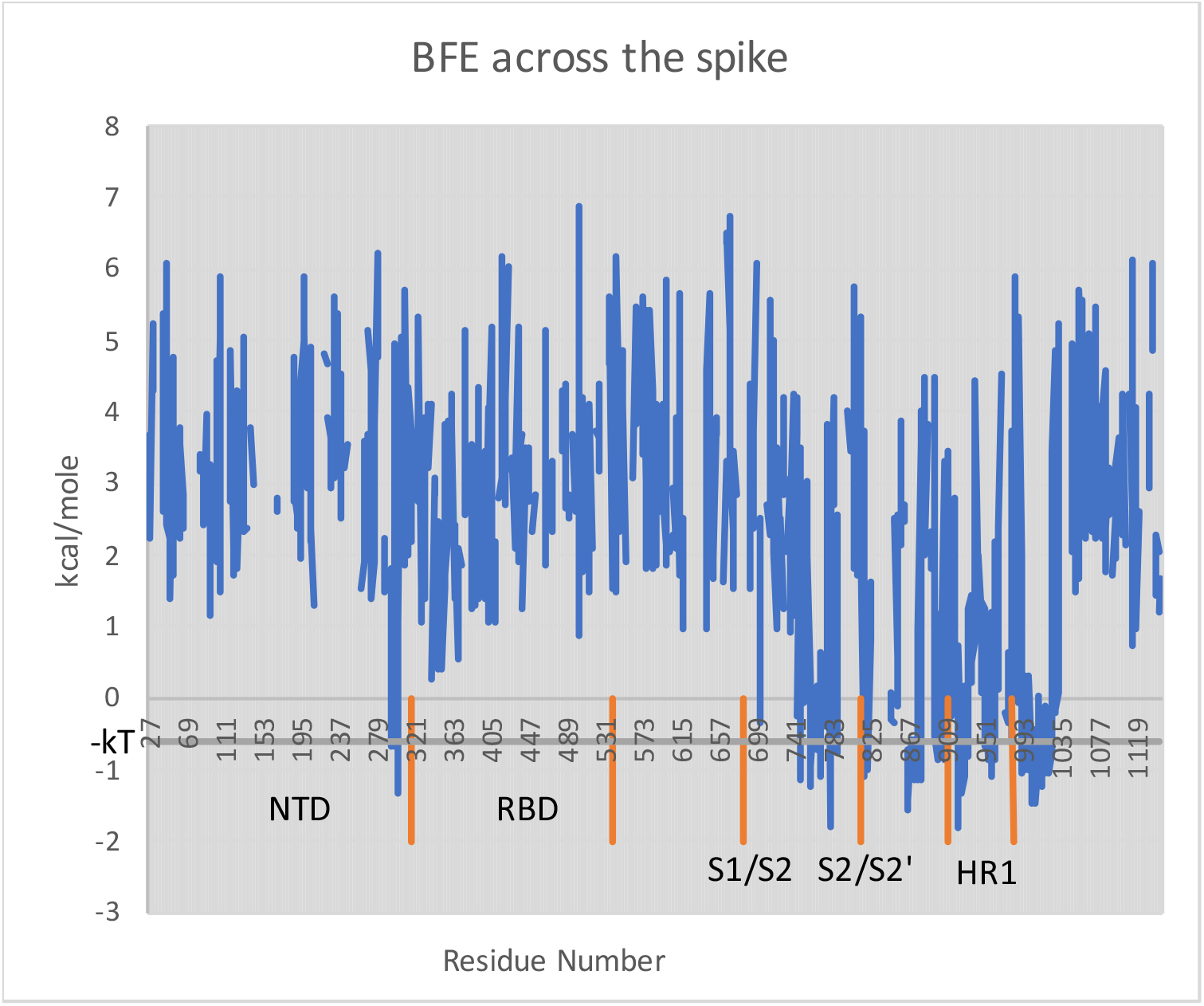
Plot of BFE by residue across the PDB-covered spike, residues 27-1147 of S. Illustrated in orange are the respective residue ranges for the N-Terminal Domain, the Receptor Binding Domain, the S_1_/S_2_ cleavage, the 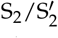 cleavage, and the first Heptad Repeat Domain. The grey horizontal line indicates one “heat quantum” *kT* ≈ 0.6 kcal/mole below zero. One confirms by comparison with the structure itself that the intersections of BFE with this line correspond to *α* helices, and in fact ones whose backbone geometry is especially near ideal *α* helices according to considerations of free energy.

However as depicted in Figure 2, the single mutation D614G, which quickly globally overtook Wuhan-Hu-1 as the predominant strain, alters BFE along the entire backbone by as much at 5.10 kcal/mole at residue 134, whereas by only 0.14 kcal/mole at residue 614 itself. Thus, a single local change of primary structure can engender a long-range change of BFE across the whole spike glycoprotein.

**Figure 2.**
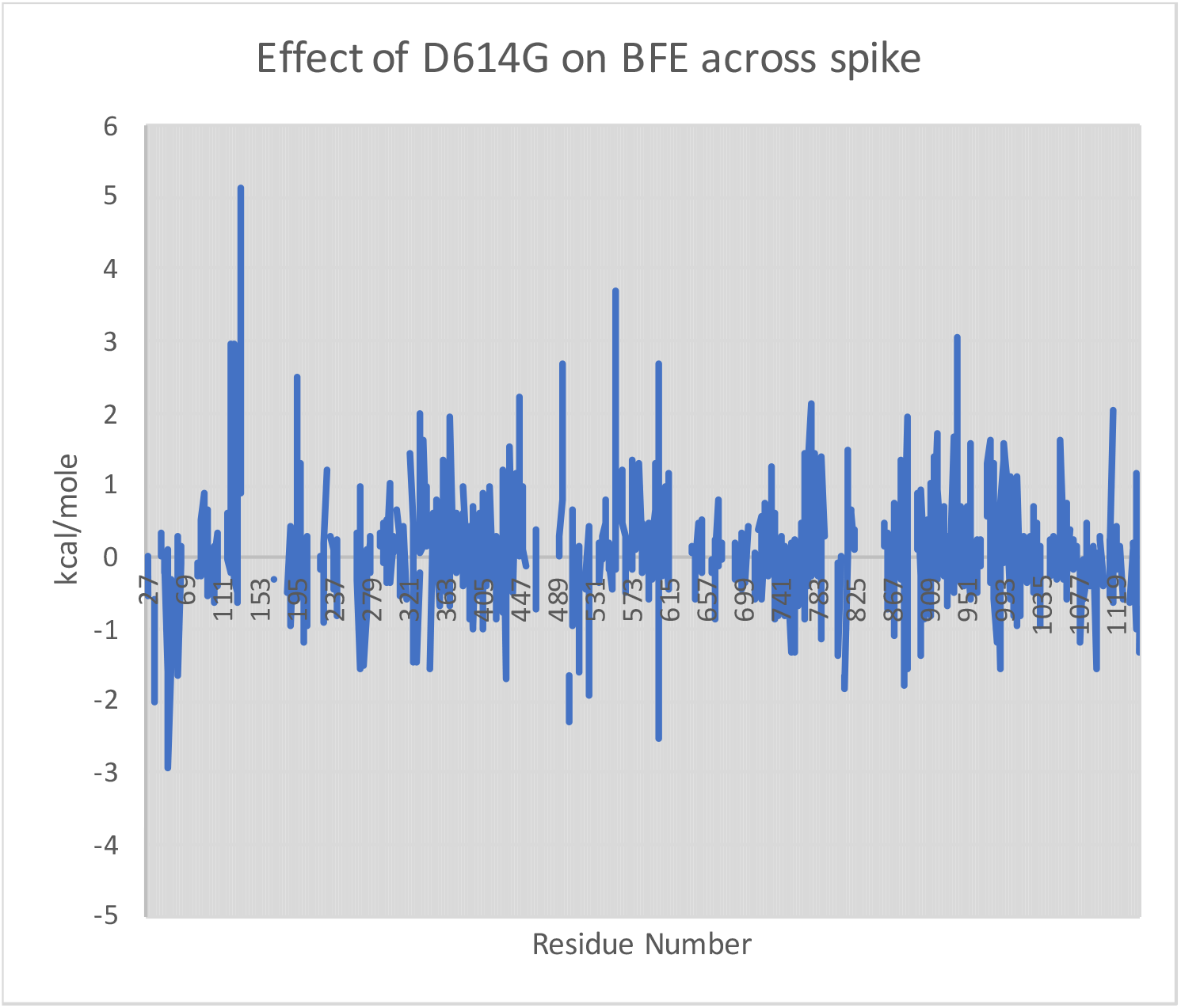
Comparison of BFE across the spike from the single mutation D614G [8]. The BFE of Wuhan-Hu-1 is computed at each residue as the average of PDB structures 7KDG and 7KDH, which are stabilized in the prefusion conformation by mutations R682G, R683G, R685G plus the 2P mutation given by K986P and V987P; the BFE of the D614G mutation is computed as the average of structures 7KDK and 7KDL, analogously stabilized but also with the D614G mutation. In each case, missing or absent residues give null. Plotted is the difference of the former minus the latter. The Wuhan-Hu-1 BFE at residue 614 itself is 2.26 kcal/mole compared to 2.12 kcal/mole for D614G, but despite this near equality at residue 614, the BFE across the entire spike is altered.

Table 1 presents findings and data about the mutating residue numbers M common to one or more of the variants under consideration here. Specifically the table summarizes BFEs and numbers of absent, missing, and unbonded residues in each of the molecules S, S_1_ and S_2_ as well as in their respective intersections 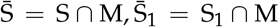 and 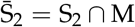 with the mutagenic residues M under consideration.

Several trends present themselves:

- S_1_ is more disorganized than S_2_ (i.e., # missing is larger);
- there are more loops in S_1_ than S_2_ (i.e., # absent is larger);
- the BFE of S_1_ is larger than S_2_;
- the same three assertions above hold for 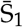 compared to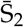 ;
- a greater ratio 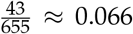 of residues in S_1_ are mutating in the variants under consideration than 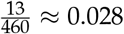 in S_2_.

Moreover, this table quantifies our main new

#### Basic Finding

The residues mutating in the variants under consideration are more often unbonded in S_1_ and bonded in S_2_.

Note that the missing and absent columns in the table come directly from the PDB and DSSP with no provisos (other than those conventions in parentheses in the text). The unbonded column presents our novel insights and depends upon a cutoff 5, namely, it is bonded (i.e., neither absent nor missing) in at least 5 of the 15 structures. The last BFE column depends not only on the cutoff 5 but also on our theory of BHBs. All of the preceding trends are invariant under changing this cutoff by unity, with this data not presented.

### 3.2. Wuhan-Hu-1 to Delta to Omicron

We shall quantify the Basic Finding for the three mutational steps from Wuhan-Hu-1 (W) to Delta (Δ), W to Omicron (*O*), and Δ to *O*. The geometry of the spike for Δ is derived from the PDB files 7V7O 7V7P…7V7V in the same manner as before for W from its exemplar structures, again with a cutoff of 5.

Table 2 quantifies our Basic Finding under various scenarios, with the W ⟼ (*) and Δ ⟼*O* mutations comparable, where (*) is the union of the variants considered in the previous section. The W ⟼ *O* transition is anomalous with its much smaller percentage of unbonded mutated residues. The explanation follows from the last column, showing the large percentages of residues that are bonded in W but not in Δ, and hence of higher mutagenic potential for their transition to *O* according to the Basic Finding.

**Table 2.**
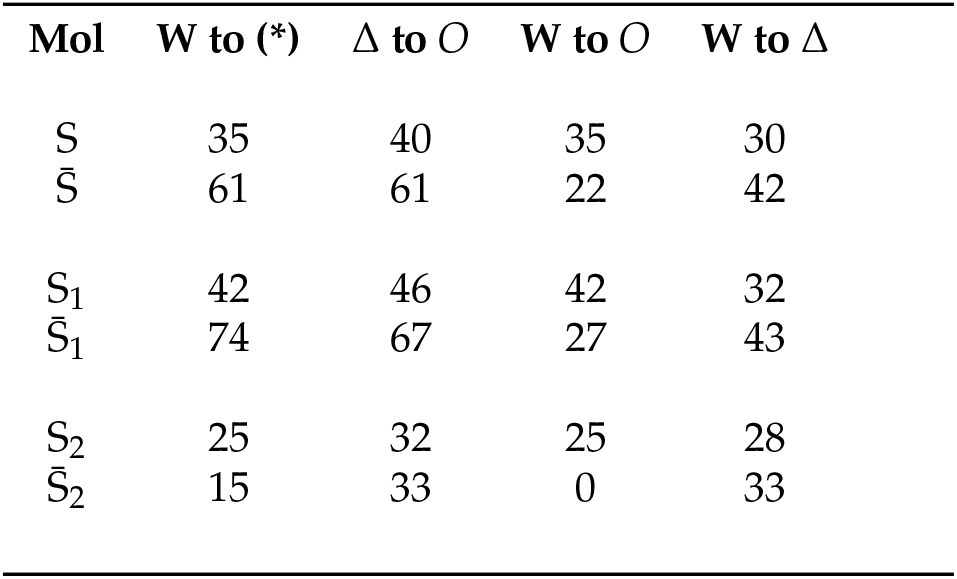
**Percentage Unbonded Residues** for each indicated molecule. W, Δ, and *O* are respectively the Wuhan-Hu-1, Delta and Omicron variants, and (*) denotes the collection of Variants of Concern or Interest or under Monitoring from [15] with mutated residues given in Table 1. It is Tulio de Oliveira’s collection of mutated residues 67 69 70 142-145 211 212 214 339 371 373 375 417 440 446 477 478 484 493 496 498 501 505 547 614 655 679 681 764 796 856 954 969 981 that serve to define *O* for our purposes here. As in Table 1, we let 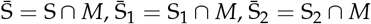 respectively denote the collection of mutagenic residues *M* in each of the molecules S,S_1_,S_2_, with *M* determined by (*) in the second column and by *O* in the third and fourth columns. The last column is simply the percentage of residues which are bonded in W and unbonded in Δ in each molecule.

The supposition that Δ played an intermediary role in the passage to *O* is already strongly bolstered by the nearly complete dominance of Δ in South Africa before the advent of *O*. Meanwhile, the PDB files 7LYK 7LYL…7LYQ [9] for the earlier South African *β* variant are of a lower quality, so their interpretation is problematic and not fully presented here; however the resulting 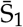 -entry for *β* to *O* in Table 2 is 48%.

## 4. Discussion

We find that mutagenic pressure on S_1_ exceeds that on S_2_, as expected based on function and location of both subunits, and that the former is more disorganized and with a lower percentage of bonded residues than the latter. These latter findings are consistent with the general trend that B-factors [3] in the receptor-binding subunit usually exceed those in the fusion subunit of a viral glycoprotein, at least in the prefusion conformation. It is argued that mutation of unbonded residues avoids the twofold constraint on

BFE imposed by metastability of the viral glycoprotein, thus explaining the tendency of mutating residues in S_1_ to be unbonded. However, among the mutating residues (19) 156-158 452 478 614 681 950 defining Delta, only 614 and 950 are bonded in Wuhan-Hu-1, in keeping with the Basic Finding, while only 681 is **unbonded** in Delta. This is fascinating and shows that there is more to our energetic argument than simply mutations avoiding BHBs, rather: Wuhan-Hu-1 to Delta is not reversible, and there is a thus an evolutionary dynamics of fixing BHBs for function and erasing them to enhance mutation in light of the Basic Finding, at least in this case of Wuhan-Hu-1 to Delta. In contrast, among the mutated residues 80 215 417 484 501 614 701 defining Beta, only 417 614 and 701 are bonded in Wuhan-Hu-1, while these plus 484 are bonded in Beta, here using any cutoff greater than unity, so bonded mutagenic residues remain bonded in this case. Backbone hydrogen bonds therefore provide an additional level of regulation of viral mutation, and this warrants further study.

As was already mentioned, the different pH of activation for the two subunits S_1_ and S_2_ may explain the opposite trend in the latter that mutating residues tend to be bonded, since the prefusion stabilized spike structures may better reflect the actual geometry and consequent BHBs of S_1_ compared to S_2_. Another related possibility is that pre-cleavage, S_1_ sits on top of S_2_ as a kind of cap, thereby sterically constraining the latter, so the active geometry of S_2_ is displayed only post-cleavage and in the course of acidic post-fusion reconformation.

In any case, the findings on S_1_ suggest a strategy for anticipating residues primed for mutation therein. However going forward, it is the residues that are unbonded for the currently mutated variants, rather than for Wuhan-Hu-1, that should be considered as likely future candidates, just as in our analysis of Delta to Omicron.

## 5. Conclusion

Our findings admit explanation by general principles, and so may hold more generally, to wit: being free from backbone hydrogen bonds increases the mutagenic potential within the receptor-binding subunit of a viral glycoprotein, and therefore deleting backbone hydrogen bonds, within the constraints of molecular functionality, can increase the mutagenic potential of a glycoprotein.

These considerations may be of utility for anticipating mutagenic pressure within the receptor-binding subunit of a viral glycoprotein based upon a lack of backbone hydrogen bonds, and this may be of substance in general for vaccine design.

## Acknowledgments

It is a pleasure to thank Pablo Guardado-Calvo, Minus van Baalen, Charles Swerdlow and the referees for critical comments.

## Abbreviations

The following abbreviations are used in this manuscript:

BFE: Backbone Free Energy
BHB: Backbone Hydrogen Bond
PDB: Protein Data Bank
W: Wuhan-Hu-1 SARS-CoV-2 strain
*β*,Δ,*O*: Beta, Delta, Omicron SARS-CoV-2 Variant

